# Lasting effect of psilocybin on sociability can be blocked by DNA methyltransferase inhibition

**DOI:** 10.1101/2025.03.10.642385

**Authors:** Chenchen Song, Tinya Chang, Tobias Buchborn, Thomas Knöpfel

**Affiliations:** Lee Kong Chian School of Medicine, Nanyang Technological University, Singapore; Laboratory for Neuronal Circuit Dynamics, Imperial College London, London, UK; Institute of Psychopharmacology, Central Institute of Mental Health, Medical Faculty Mannheim, University of Heidelberg, Heidelberg, Germany; Laboratory for Neuronal Circuit Dynamics, Hong Kong Baptist University, Hong Kong SAR

## Abstract

The recent renaissance in research on psychedelics such as psilocybin has highlighted their therapeutic potential including their lasting influences on brain function. Here we report that a single systemic administration of the serotonergic psychedelic psilocybin can durably promote social behaviour in the Cntnap2-knockout mouse model of autism. This effect can be blocked by pharmacological inhibition of DNA methyltransferase I, indicating an epigenetic mechanism underlying the long-lasting effect of psilocybin.

## Introduction

The serotonergic psychedelic psilocybin has been shown to induce behavioural alterations outlasting the acute psychedelic effect. Clinical studies have revealed a therapeutic potential of psilocybin for the treatment of psychiatric disorders^1^ and suggested beneficial effects on behavioural traits associated with and common to several neuropsychiatric conditions^2^. Intriguing but mechanistically unexplained are the therapeutic effects that last at least several weeks, for instance the alleviation of depression in treatment-resistant patients following a single dose of psilocybin-assisted treatment^3–5^. Reduced sociability is a common characteristic of all types of autism under the classification autism spectrum disorder (ASD) and this altered trait is replicated in several mouse models that mimic the genetics of ASD^6–8^. Here, we examined the effect of a single systemic injection of psilocybin in altering the social behaviour of the Cntnap2 knockout mouse model^8^ of autism spectrum disorder and the durability of this effect.

## Results

### Psilocybin durably restores sociability in Cntnap2^-/-^ homozygous mutant mice

We administered a single systemic injection of 1 mg/kg psilocybin or saline to adult Cntnap2 homozygous mutant (Cntnap2^-/-^ KO) and Cntnap2 homozygous wild-type isogenic control (Cntnap2^+/+^ WT) mice at 8 weeks of age. We assessed the social behaviour of these mice using the three-chamber social behaviour test at 1 day, 1 week, and 2 weeks after the single injection, corresponding to ages of postnatal week 8, 9 and 10 respectively. We quantified sociability as the amount of time a mouse approaches and explores a cup containing another mouse versus an empty cup. Similar to the reduced sociability shown in several previous studies^8–10^, saline treated homozygous mutant KO mice exhibited a reduced sociability compared to the isogenic wild-type control group (p = 0.042, Cntnap2^-/-^*_Vehicle_* vs Cntnap2^+/+^ *_Vehicle_*, Bonferroni’s post-hoc). Importantly, however, psilocybin-treated homozygous mutant KO group showed a significant increase in sociability compared to the vehicle-treated Cntnap2^-/-^_Vehicle_ KO control group (p < 0.001, Cntnap2^-/-^*_Psilocybin_* vs Cntnap2^-/-^*_Vehicle_*, Bonferroni’s post-hoc, **Fig 1a**) and this effect was sustained for at least two weeks following the single psilocybin administration. Sociability of Cntnap2^-/-^*_Psilocybin_* KO group was restored to a level similar to the isogenic wild-type control group (p = 0.796, Cntnap2^-/-^*_Psilocybin_*vs Cntnap2^+/+^*_Vehicla_*, Bonferroni’s post-hoc). The total time a mouse spent exploring any cup (either mouse-containing or empty) did not significantly differ between treatment of the same genotype groups (p > 0.999, Cntnap2^-/-^*_Vehicle:Vehicle_* vs Cntnap2^-/-^*_DNMTI:Psilocybin_*; p = 0.390, Cntnap2^+/+^*_Psilocybin_* vs Cntnap2^+/+^*_Vehicle_* Bonferroni’s post-hoc; **Extended Data Fig 1a**), indicating that the difference in sociability is due to a specific deficit in social behaviour rather than a variability in the drive to explore. We did not detect any significant effect of sex on our observations (p = 0.850, repeated measure ANOVA with Bonferroni’s post-hoc).

**Figure 1.**
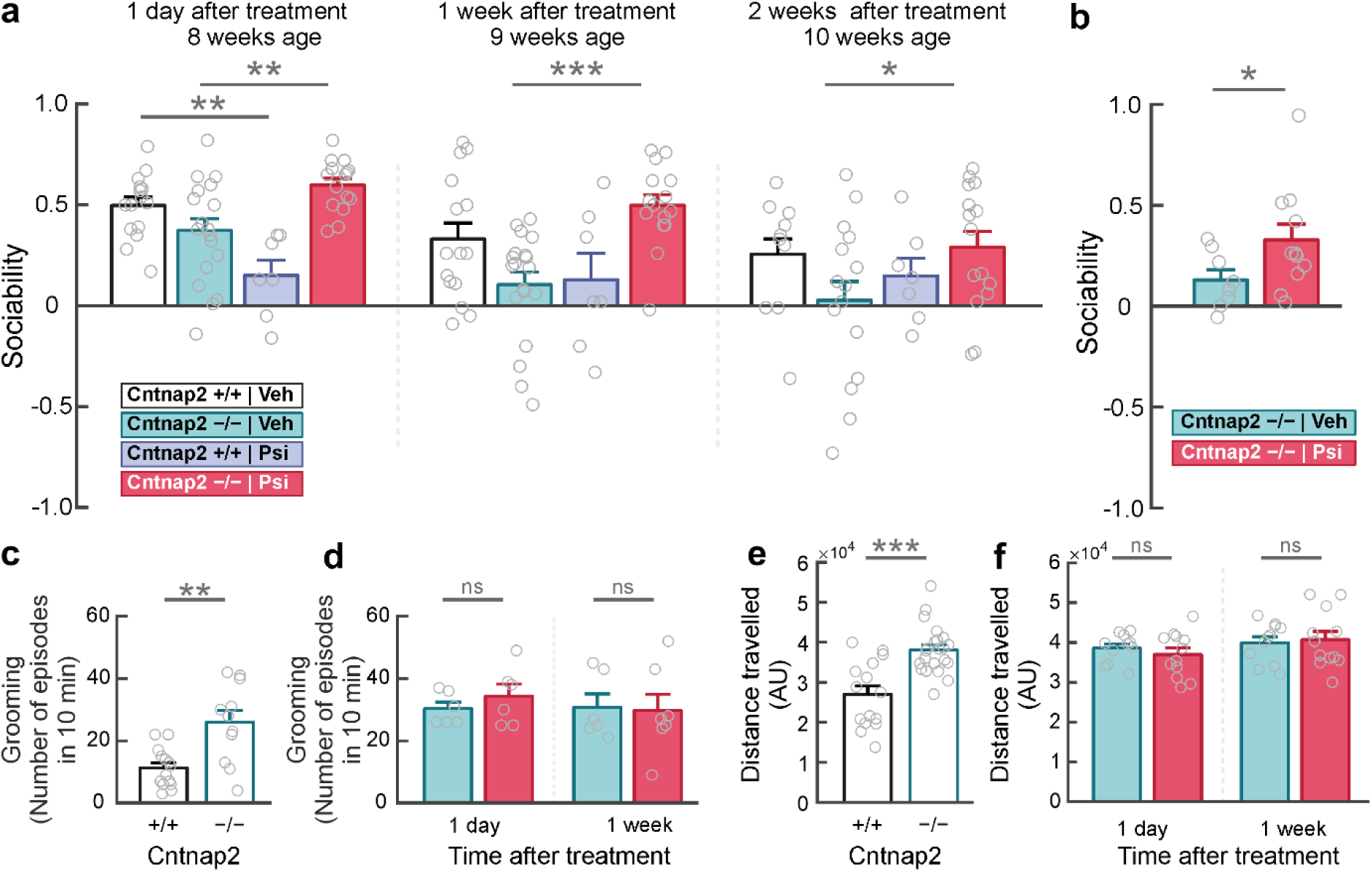
Psilocybin durably restores sociability in Cntnap2^-/-^ homozygous KO mice. **a.** A single injection of psilocybin rescues the social deficits in Cntnap2^-/-^ KO mice for up to two weeks (**Extended Tables 1a, 2**). **b.** Psilocybin’s sociability rescuing effect is observed in Cntnap2^-/-^ KO mice tested for the first time at two weeks after a single treatment. **c-d.** The significantly higher repetitive behaviour displayed by Cntnap2^-/-^ KO mice (**c**) is not altered by psilocybin (**d**). **e-f.** The significant hyperactivity displayed by Cntnap2^-/-^ KO mice (**e**) is not altered by psilocybin (**f**). Mean ± SEM shown. Individual data point shown. For detailed statistics see **Extended Data Table 1-3**. *p < 0.05, ** p < 0.01, ***p < 0.001.

Further, we treated a subset of Cntnap2^-/-^ KO mice with a single administration of psilocybin or saline vehicle, and tested for their sociability for the first time at two weeks after treatment. Here we also observed significantly higher sociability in psilocybin treated Cntnap2^-/-^ KO mice (p = 0.025, Cntnap2^-/-^*_Psilocybin_* vs Cntnap2^-/-^*_Vehicle_*; **Fig 1b**, **Extended Data Table 2a**). This observation indicated that our observation of psilocybin effect was indeed a sustained rescue of social behaviour rather than an effect of a repeated testing design. Given this observation, we next sought to examine each timepoints of our repeated measure observation, and detected the greatest psilocybin rescuing effect at 1 week post-administration (at 1 day: p = 0.007, at 1 week: p < 0.001, at 2 weeks: p = 0.028, Cntnap2^-/-^*_Psilocybin_* vs Cntnap2^-/-^*_Vechicle_*, **Fig 1a**, **Extended Data Table 2b**).

### No detectable effects of psilocybin on two other behavioural measures

Then, we explored whether psilocybin also affected other aspects of behaviour that are not directedly related to sociability but are possibly affected in ASD model mice^8,11^. We examined repetitive behaviour by quantifying self-grooming activity during habituation. We observed that Cntnap2^-/-^ homozygous mutant KO mice displayed significantly higher number of self-grooming episodes compared to Cntnap2^+/+^ homozygous WT mice during the final habituation session prior to drug treatments (p = 0.002, Cntnap2^+/+^ vs Cntnap2^-/-^, Mann Whitney, **Fig 1c**). No significant difference was detected in the number of self-grooming episodes displayed by Cntnap2^-/-^ KO mice at 1 day after psilocybin treatment (p = 0.329, Cntnap2^-/-^*_Vehicle_* vs Cntnap2^-/-^*_Psilocybin_*, Mann Whitney, **Fig 1d**) or at 1 week after drug treatment (p = 0.461, Cntnap2^-/-^*_Vehicle_* vs Cntnap2^-/-^*_Psilocybin_*, Mann Whitney, **Fig 1d**). We therefore concluded that repetitive behaviour, although increased in Cntnap2^-/-^ KO mice, is not affected by psilocybin.

Further, we tracked the positions of the test mice during the habituation phase without the introduction of a social partner. In line with previous reports of hyperactivity in Cntnap2^-/-^ KO mice^8^, we observed that Cntnap2^-/-^ KO mice displayed significantly higher distance travelled during the 10 minutes observation period compared to wildtype (p = 0.001, Cntnap2^+/+^ vs Cntnap2^-/-^, Dunn’s post-hoc, **Fig 1e**). Despite drug treatment, no significant difference was detected in the distance travelled by Cntnap2^-/-^ KO mice at 1 day after drug treatment (p = 0.518, Cntnap2^-/-^ vs Cntnap2^-/-^, Mann Whitney) or at 1 week after drug treatment (p = 0.976, Cntnap2^-/-^ vs Cntnap2^-/-^, Mann Whitney). We therefore concluded that psilocybin has no lasting effect on spontaneous locomotory drive.

### Psilocybin does not enhance social behaviour in the Cntnap2 genotypes without social deficit

Interestingly, the single administration of psilocybin reduced sociability in Cntnap2^+/+^ isogenic wild-type group, at 1 day post-treatment (p = 0.001, Cntnap2^+/+^_Vehicle_ vs Cntnap2^+/+^_Psilocybin_). Psilocybin-treated isogenic wild-type mice remained lower in sociability scores at 1 week (p = 0.155, Cntnap2^+/+^_Vehicle_ vs Cntnap2^+/+^_Psilocybin_) and at 2 weeks (p = 0.207, Cntnap2^+/+^_Vehicle_ vs Cntnap2^+/+^_Psilocybin_) after treatment although no longer statistically significantly (**Fig 1a**). In line with the idea that *CNTNAP2* is a recessive gene^12^, heterozygous Cntnap2^+/-^_Vehicle_ group did not show significantly altered sociability compared to wildtype Cntnap2^+/+^_Vehicle_ group (p = 1.000, Cntnap2^+/-^_Vehicle_ vs Cntnap2^+/+^_Vehicle_, Bonferroni’s post-hoc, **Extended Data Table 1b**). Cntnap2^+/-^ heterozygous mice also displayed reduced sociability after psilocybin treatment (**Extended Data Fig 1b**), even though not statistically significantly. These observations indicate that while the prosocial effect of psilocybin has potential to rescue abnormal behavioural traits, in the absence of behavioural deficits psilocybin does not enhance and may even reduce sociability.

### Lack of correlation between sociability and the acute effect of psilocybin

Psilocybin or its active metabolite psilocin, like other psychedelic 5HT2AR agonists, trigger an acute head twitch response (HTR)^13^ that lasts for the duration of the agonist’s bioavailability^14^. Immediately after psilocybin application, we assayed the HTR for a 20 minutes observation period to confirm the normal functioning of 5HT2ARs in psilocybin-treated mice (**Extended Data Fig 1c**, **Extended Data Table 4a**). No difference in HTR was observed between the different genotype groups without psilocybin (p = 0.825, Cntnap2^+/+^_Vehicle_ vs Cntnap2^-/-^_Vehicle_; p > 0.999, Cntnap2^+/+^_Vehicle_ vs Cntnap2^+/-^_Vehicle_, Dunn’s post-hoc), and psilocybin significantly increased HTR in all genotype groups (p < 0.001, Cntnap2^+/+^_Vehicle_ vs Cntnap2^+/+^_Psilocybin_; p = 0.001, Cntnap2^-/-^_Vehicle_ vs Cntnap2^-/-^_Psilocybin_; p = 0.006, Cntnap2^+/-^_Vehicle_ vs Cntnap2^+/-^_Psilocybin_ Dunn’s post-hoc) to a similar level (p > 0.9999, Cntnap2^+/+^_Psilocybin_ vs Cntnap2^-/-^_Psilocybin_; p > 0.9999, Cntnap2^+/+^_Psilocybin_ vs Cntnap2^+/-^_Psilocybin_; Dunn’s post-hoc). Even though sex-related differences in HTR have been reported for other psychedelic compounds and analoqgues^15,16^, we did not detect any significant effect of sex on the HTR (p = 0.452, Cntnap2^-/-^_Psilocybin_:Male vs Cntnap2^-/-^_Psilocybin_:Female, Mann Whitney; **Extended Data Table 4b**) which is similar to a lack of sex-specific effects on psilocybin-induced HTR as observed by others^17^. Moreover, we observed no significant correlation between acute HTR and sociability levels in the Cntnap2^-/-^ KO mice (**Extended Data Table 4c**). This indicates that the reduced sociability in Cntnap2^-/-^ KO mice cannot be explained by an altered acute function of 5HT2ARs.

### DNA methyltransferase inhibition blocks psilocybin-induced sociability changes in Cntnap2^-/-^ homozygous mutant KO mice to a similar extent as 5HT2AR antagonism

Given that the observed sociability-rescuing effect outlasts the bioavailability of psilocybin, we reasoned that the lasting alterations in sociability following psilocybin administration may originate from mechanisms other than sustained 5HT2AR activation^18^. We reasoned that epigenetic alterations may provide a potential mechanism for the observed lasting social deficit rescuing effects of psilocybin. We focused on changes in DNA methylation in view of reports showing differential DNA methylation patterns in autism spectrum disorder^19,20^.

We pretreated mice with systemic administration of RG108^21–23^ - a small-molecule inhibitor of DNA methyltransferase I (DNMT1) - to test whether DNMT inhibition (DNMTi) prevents the effects from psilocybin administration. We then behaviourally tested the mice for sociability at 1 day and 1 week after treatment (**Fig. 2**). When pretreated with vehicle, psilocybin treatment rescues the reduced sociability in homozygous KO mice (p < 0.001, Cntnap2^-/-^_Vehicle::Vehicle_ vs Cntnap2^-/-^_Vehicle::Psilocybin_, Bonferroni’s post-hoc), similar to our earlier observations. DNMTi pretreatment significantly reduced the psilocybin-induced increased sociability in Cntnap2^-/-^ KO mice (p = 0.003, Cntnap2^-/-^_Vehicle::Psilocybin_ vs Cntnap2^-/-^_DNMTi::Psilocybin_, Bonferroni’s post-hoc) and psilocybin treatment is no longer effective (p = 1.000, Cntnap2^-/-^DNMTi::Psilocybin vs Cntnap2^-/-^DNMTi::Vehicle, Bonferroni’s post-hoc). DNMTi pretreatment is similarly effective at 1 day and 1 week after administration (**Extended Data Table 6**). This indicates that *de novo* DNA methylation is required for the sustained effect of psilocybin on sociability in Cntnap2^-/-^ KO mice. DNMTi pretreatment itself did not affect sociability in Cntnap2^-/-^ KO mice (p = 1.000, Cntnap2^-/-^_Vehicle::Vehicle_ vs Cntnap2^-/-^_DNMTi::Vehicle_, Bonferroni’s post-hoc), or in Cntnap2^+/+^ isogenic WT mice (p = 0.526, Cntnap2^+/+^_Vehicle::Vehicle_ vs Cntnap2^+/+^_DNMTi::Vehicle_, Repeated measure ANOVA). Additionally, we confirmed that DNMTi pretreatment in isogenic WT mice (**Extended Data Table 7**) does not have an effect on the number of self-grooming episodes (p = 0.222, Cntnap2^+/+^_Vehicle::Vehicle_ vs Cntnap2^+/+^_DNMTi::Vehicle_, Mann Whitney), or total distance travelled (p = 0.128, Cntnap2^+/+^_Vehicle::Vehicle_ vs Cntnap2^+/+^_DNMTi::Vehicle_, Mann Whitney).

**Figure 2.**
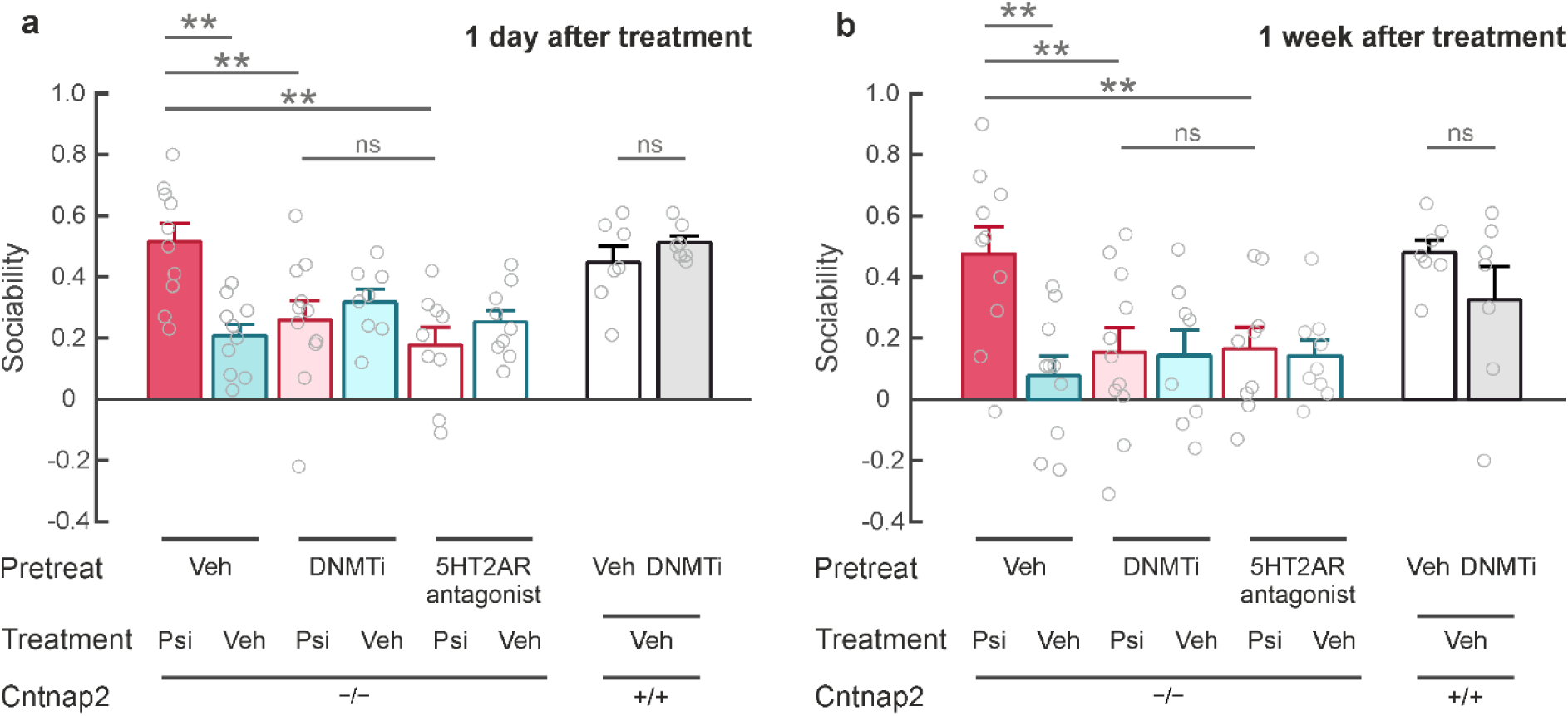
Psilocybin’s sociability rescuing effect is blocked by DNMTi. **a.** Psilocybin’s sociability rescuing effect can be blocked by pretreatment of DNMTi to a similar extent as 5HT2AR antagonism. **b.** The pretreatment blockade effects are sustained at 1 week after treatment. Mean ± SEM shown. Individual data point shown. For detailed statistics see **Extended Data Table 5-6**. ** p < 0.01.

Psilocybin is known to act predominantly through the activation of serotonin 2A receptors (5HT2ARs)^24–26^. To test whether the observed effects of psilocybin were indeed initiated by 5HT2AR activation, we also performed similar experiments using pretreatment with a specific 5HT2AR antagonist volinanserin (MDL100907). We observed that 5HT2AR antagonist pretreatment prevented the psilocybin-induced increased sociability in Cntnap2^-/-^ KO mice (p = 0.001, Cntnap2^-/-^_Vehicle_:Psilocybin vs Cntnap2^-/-^*_5HT2AR-ant::Psilocybin_* groups, Bonferroni’s post-hoc; **Fig 2**, **Extended Data Table 5**). When comparing mice pretreated with DNMTi or 5HT2AR antagonist, we found no significant difference in psilocybin’s effect on sociability (p = 1.000, Cntnap2^-/-^*_DNMTi::Psilocybin_* vs Cntnap2^-/-^*_5HT2AR-ant::Psilocybin_*groups, Bonferroni’s post-hoc; **Fig 2**, **Extended Data Table 5**), which indicates that DNMTi is as effective as 5HT2AR antagonist in blocking psilocybin’s effect on rescuing sociability. Further, we observed that the blocking effect of both pretreatment groups were sustained at 1 week after treatment (**Fig 2b**; **Extended Data Table 6**). This indicated that both *de novo* DNA methylation and 5HT2AR activation are required for the sustained effect of psilocybin on sociability in Cntnap2^-/-^ KO mice.

While pretreatment with 5HT2AR antagonist effectively abolished the psilocybin-induced head-twitch response (p < 0.001, Cntnap2^-/-^_Vehicle::Psilocybin_ vs Cntnap2^-/-^_5HT2AR-ant::Psilocybin_ groups, Dunn’s post-hoc), psilocybin-induced HTR was not affected by pretreatment with DNMTi (p > 0.999, Cntnap2^-/-^_Vehicle::Psilocybin_ vs Cntnap2^-/-^_DNMTi::Psilocybin_ groups, Dunn’s post-hoc) or with vehicle (p > 0.999, Cntnap2^-/-^_Vehicle::Psilocybin_ vs Cntnap2^-/-^DNMTi::Psilocybin groups, Dunn’s post-hoc; **Extended Data Fig 2**, **Extended Data Table 8**). These observations indicate that the effect of pretreatment with DNMTi did not interfere with the normal acute effect of 5HT2AR activation by psilocybin, or spontaneous head-twitch events in Cntnap2^-/-^ KO mice (p > 0.999, Cntnap2^-/-^_Vehicle::Vehicle_ vs Cntnap2^-/-^_DNMTi::Vehicle_ groups, Dunn’s post-hoc; **Extended Data Fig 2**, **Extended Data Table 8**).

## Discussion

Taken together, our observations demonstrate that psilocybin rescues social deficits in the Cntnap2 knock-out mouse model of altered sociability in ASD, and this effect is sustained for at least up to two weeks. Our results further suggest that the long-lasting effect of psilocybin depends on *de novo* DNA methylation dynamics, since pretreatment of DNA methyltransferase inhibitor is as effective in blocking psilocybin’s therapeutic effect as antagonism of psilocin’s main binding target 5HT2A receptor. Our work here supports the rapidly expanding body of experimental evidence that psychedelics have the capability to exert sustained effects on brain function and behaviour, and an involvement of epigenetic mechanisms in this durable effect.

Despite this promising therapeutic effect, we also note that psilocybin does not improve sociability in mice that do not show behavioural deficits, such as that of the Cntnap2 isogenic WT or heterozygote groups here. Our observations even indicate that psilocybin exerts opposing effects, reducing sociability in the absence of abnormal sociability traits. Thus, it appears that psilocybin is not a general sociability enhancer, but rather is only beneficial when in deficit, a clinically relevant behavioural measure for brain disorders or mental health issues. When viewing psilocybin administration as a pharmacological perturbation, its counteracting effects suggest the existence of a setpoint that governs social behaviour. Our observation that DNA methylation plays a role in psilocybin’s sustained social deficit rescuing effect further suggests that the setpoint for social behaviour may be controlled and/or reconfigured through epigenetic plasticity mechanisms. In line with this, others have observed that a single administration of other serotonergic psychedelic such as lysergic acid diethylamide (LSD) into control mice also did not enhance social behaviour^27^. However, social enhancement could be achieved through repeated LSD treatments^27^, and repeated LSD treatments into control mice has been separately reported to result in detectable changes in DNA methylation in the brain^28^. Considering this together with our observations, these experimental evidence collectively suggest that different behavioural setpoints may exist for normal vs deficit states, which may be adjusted with varying levels of ease.

Separately, the single administration of another serotonergic psychedelic 1-(2,5-dimethoxy-4-iodophenyl)-2-aminopropane (DOI) led to detectable changes in chromatin organisation and enhancer activity, associated with increased synaptogenesis^29^. Psilocybin may similarly lead to epigenomic changes including alteration of DNA methylation dynamics, which may underpin our observations here. In the case of the socially reduced Cntnap2^-/-^ homozygous mutant KO mice, we observed that pretreatment with DNMTi is not the exclusive mechanism to block psilocybin’s therapeutic effects, but it is as effective as pretreatment with 5HT2AR antagonism. This suggests that psilocybin-induced changes in DNA methylation dynamics is downstream of 5HT2AR activation. Future work will be necessary to identify the signalling pathways involved.

Further, we noted a general decline in sociability across postnatal weeks 8-10 for all experimental groups in line with previously reported age-dependent reduction in social-related behaviour in mice^30^. This decline is much more significant in Cntnap2 KO mice (p = 0.003, Cntnap2^-/-^_Vehicle_:1 day vs Cntnap2^-/-^ vehicle:1 week; p = 0.532, Cntnap2^-/-^ vehicle: 1 week vs Cntnap2^-/-^ vehicle: _2 weeks_; p = 0.008, Cntnap2^-/-^ vehicle:1 day vs Cntnap2^-/-^ vehicle: 2 weeks; T test) and is delayed following psilocybin treatment (p = 0.120, Cntnap2^-/-^_Psilocybin_:1-day vs Cntnap2^-/-^_Psilocybin_: 1 week; p = 0.040, Cntnap2^-/-^_Psilocybin_: 1 week vs Cntnap2^-/-^_Psilocybin_: 2 weeks; p = 0.002, Cntnap2^-/-^_Psilocybin_:1 day vs Cntnap2^-/-^_psilocybin_: ^2^^weeks^; T test). It will be interesting to investigate whether psilocybin extends or even re-opens the critical developmental window where experience-dependent components of sociability traits are the most plastic, and indeed, latest experimental observations by others indicate that psychedelics has the potential to re-open the critical period for social reward learning^31^. This appears to be in line with our observations here that while psilocybin was effective in rescuing social deficits, it did not alter the increased self-grooming repetitive behaviour or hyperactivity associated with this ASD model^8^, which may not involve critical periods.

## Methods

### Animals

Experimental procedures at Nanyang Technological University Singapore were performed in accordance with approved protocols following review by the Institutional Animals Care and Use Committee. Experimental procedures performed at Imperial College London UK in accordance with the United Kingdom Animal Scientific Procedures Act (1986), under Home Office Personal and Project Licences following appropriate ethical review. Cntnap2^-/-^ homozygous mutant KO^8^ transgenic mice (JAX017482, Jackson Labs USA) were maintained on a C57bl6/J background. Offsprings were weaned at P21 and housed in groups of up to five animals per cage after weaning. All animals were maintained in individually ventilated cages, on a 12/12 h light/dark cycle (light on between 07:00-19:00) at 21 ± 2°C and 55 ± 10% humidity. Water and food were provided ad libitum. 8-10 weeks old mice of both genders were used for behavioural experiments. Mice were group-housed where possible, with the same littermate mice both before and after treatments. Mice in the same cage typically included treatments of different conditions and/or included untreated companion mice. All experiments were conducted during the light phase.

### Drug administration

Psilocybin (Usona Institute US) was dissolved in sterile saline solution (0.9% NaCl) to appropriate concentrations. 8-week old offspring were intraperitoneally injected either with saline or psilocybin (1 mg/kg) 24 hours prior to behavioural testing. We chose this dose of psilocybin based on our previous observations^32^ as well as observations from several other groups^17,33^. For the duration of the acute drug effect, each mouse was placed in a separate chamber to assess the head-twitch response.

### Head-twitch response

Head-twitch events were evaluated immediately after intraperitoneal administration of either saline or psilocybin for a period of 20 minutes. The mouse was placed into a customized behavioural box immediately after injection and was free to explore. The number of head-twitch events were counted by direct observation in 5-minute bins. The box is of size 40 cm X 20 cm X 20 cm [L x W x H] and the floor was covered with clean bedding. Experimenters were not blind to the treatment for head-twitch counts.

### DNA methyltransferase I inhibition

We used the small-molecule DNMT1 inhibitor RG108 (Tocris, UK) to inhibit DNA methyltransferase I in vivo^21–23^. RG108 was dissolved in 10% DMSO then diluted using saline (final DMSO received by the mice was 0.75%). Two doses of 10 mg/kg RG108 or vehicle was systemically administered via intraperitoneal injections per test animal, first dose at 60 min prior to psilocybin or saline treatment, second dose together with psilocybin or saline treatment. We selected RG108 due to its low cytotoxicity and its selective binding to DNA methyltransferase I.

### 5HT2A receptor antagonism

We used the selective 5HT2A receptor antagonist MDL100907 (Tocris, UK) to inhibit 5HT2A receptor activity in vivo^34^. MDL100907 was dissolved in 10% DMSO then diluted using saline. One dose of 1 mg/kg MDL100907 or vehicle was systemically administered via intraperitoneal injections per test animal at 15 min prior to psilocybin or saline treatment.

### Behavioural testing

Social behaviour was tested in a custom-built Plexiglass three-chamber box with matt white acrylic floor. The total arena (60 x 40 x 22 cm [L x W x H]) was divided in three smaller evenly sized chambers (20 x 40 x 22 cm [L x W x H]) interconnected by 4 x 4 cm doors. Transparent cylinders (10.5 cm internal diameter, 11 cm external diameter, 16 cm length) with 1-cm slits were used as cups. The box was placed in a darkened and quite room, illuminated from above with infrared LEDs located 1 meter over the arena.

Age-matched, sex-matched non-littermate mice with no social behavioural deficits (i.e. no homozygous mutant KO mice) from the same genetic background were used as social partners.

Animals were acclimatised to the testing room for at least one hour before the start of experiment on each day. Behavioural testing took place over 4 consecutive days in the first week. Each test mouse was placed into the centre chamber and allowed to freely explore and habituate to the three-chamber setup for 10 minutes over 3 consecutive days, doors to the side chambers were kept open. Social partner mice were individually pre-habituated to the cups also for 10 minutes over 3 consecutive days. On day 4, each test animal was allowed to freely explore all three empty chambers for 10 minutes, then driven to the central chamber with the doors closed while an age-, size-, and gender-matched unfamiliar non-littermate mouse (S) was placed into a cup in one side chamber, and an empty cup (C) was placed in the opposite side chamber. The side of the social partner chamber was randomised. The test mouse was then allowed to freely explore chambers for 10 minutes. Re-testing sessions at day 7 and day 14 after administration were performed using the day 4 protocol, with new stranger mice that the test mouse never encountered previously. Sex of the test mice were randomised where possible.

All behavioural sessions were video-recorded using a CMOS camera (Basler acA2000-165umNIR) and Basler software (Basler AG, Germany). Video recordings were used to track the position of the body of the animal in each chamber, as well as to manually time the duration of the test mouse’s social exploratory behaviour by a blinded well-trained observer. We observed that the test mice frequently spent time in a side chamber while avoiding the interaction object, therefore we manually quantified the actual interaction duration of the test mouse with the cup quantified as either sniffing or crawling on a cup that is either empty or containing another mouse. We then derived a normalised sociability index using the equation:

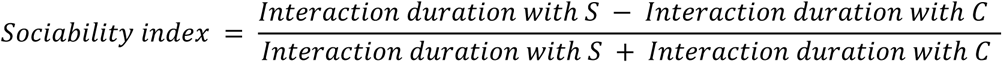

### Position tracking and analysis

We used DeepLabCut^35,36^ to track the positions of the randomly selected test mouse during the habituation phase. Each mouse was labelled by a trained network to identify at least 4 body parts including the snout, two ears, tail base. Body positions were then calculated from the label coordinates and analysed using custom codes in MATLAB. Movement speed is calculated as the difference between the body positions between two consecutive image frames.

Chamber occupancy during the habituation phase of the test were calculated. No consistent side preference were detected either for an experimental group or for the same mouse on different test sessions.

### Grooming analysis

Episodes of grooming behaviour were manually scored by an experimenter who was blind to the genotyping and treatment conditions.

### Statistical analysis

Head twitch response data were analysed using non-parametric Kruskal-Wallis test with alpha-adjusted Dunn’s multiple post-hoc comparisons, or Mann Whitney test as a priori specified. Data related to sociability index were analysed via three-factorial ANOVA with repeated measures on one factor and followed up on by Bonferroni-corrected multiple comparisons. Calculations were carried out using GraphPad Prism and SPSS, respectively. Correlations (Spearman’s rho) were calculated using MATLAB.

## Acknowledgement

We thank R Wood, B Glenn, C Day, D Macdonald, N Chua, K Lee, J Lim, Z Yong for assistance with animal husbandry. We thank MJ Hossen for technical assistance. We thank G Pearson, I Mollinedo-Gajate, T Lyons and all other members of the Knopfel lab for discussions and feedback on the manuscript. We thank CC Ng, EPY Wong, R Dienzo, MF Basir, BW Ng, P Teo, KL Lim, S Ponniah, A Tashiro, K Wong, G Augustine, W Tay, QJ Liew, YS Yip, CY Chin, J Ng, C Rantle, R Tyacke, D Nutt for administrative and infrastructure support. This work was supported by: US National Institute of Health BRAIN Initiative Grant (5U01NS099573) to TK, Hong Kong STEM Initiative Funds (Hong Kong Jockey Club Charities Trust) to TK, Research Matching Grant (Government of Hong Kong, TFD2023-P05-02) to TK, Lee Kuan Yew Postdoctoral Fellowship administered by Nanyang Technological University Singapore (022506-00001) to CS, and Open Fund Young Individual Research Grant (MOH-001720) administered by the Singapore Ministry of Health’s National Medical Research Council to CS. This work was supported by the Investigational Supply Program of Usona Institute US.

## Contributions

CS and TK designed the study. CS and TC performed the experiments. CS, TK and TB analysed the data. CS and TK supervised the project and wrote the manuscript with inputs from all authors.

## Ethics declarations and competing interests

The authors report no competing interests.

## Data availability

Data is available from the corresponding author upon reasonable request.

## Overview of Extended Data

### Extended Figures

Extended Figure 1: Parameters related to psilocybin treatment of Cntnap2 mice.

Extended Figure 2: Head-twitch response related to the pretreatment experiment.

### Extended Tables

Extended Table 1: Statistics for durability of psilocybin-induced effects on sociability, relates to Fig 1.

Extended Table 2: Statistics for durability of psilocybin-induced effects on sociability by test session, relates to Fig 1.

Extended Table 3: Statistics for repetitive behaviour and locomotion, relates to Fig 1.

Extended Table 4: Statistics for head-twitch response, and HTR-SI correlations.

Extended Table 5: Statistics for either DNMTi or 5HT2AR antagonist pretreatment on psilocybin’s sociability effect, relates to Fig 2.

Extended Table 6: Statistics for either DNMTi or 5HT2AR antagonist pretreatment on psilocybin’s sociability effect by test session, relates to Fig 2.

Extended Table 7: Statistics for the effects of DNMTi on the behaviour of isogenic wild-type mice.

Extended Table 8: Statistics for either DNMTi or 5HT2AR antagonist pretreatment on psilocybin’s effect on HTR.

---

**Extended Data Figure 1.**
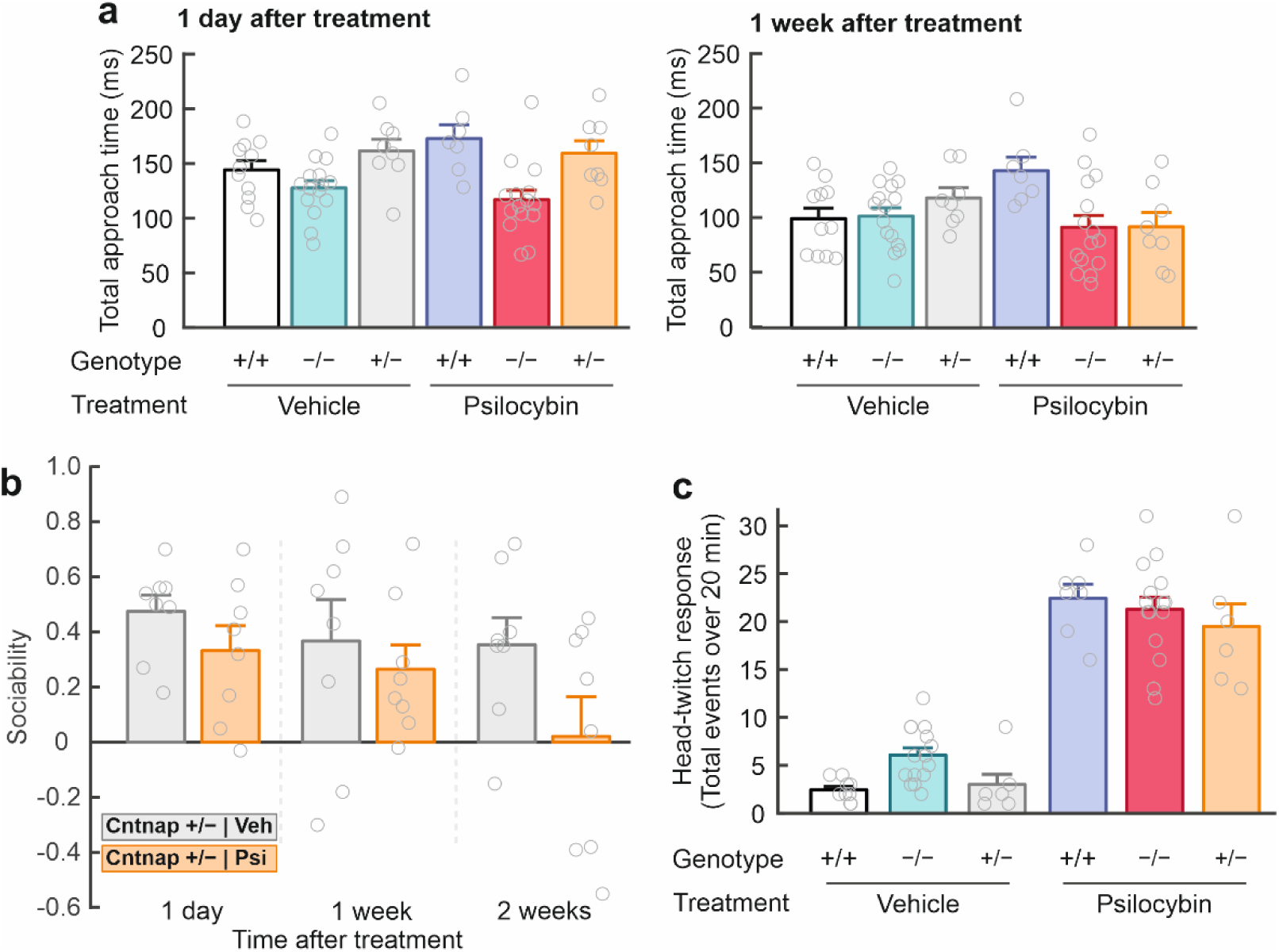
**a.** Total approach time for at 1 day (left) and 1 week (right) after treatment. **b.** Sociability at 1 day, 1 week and 2 weeks post-treatment (1 mg/kg psilocybin or vehicle control) for Cntnap2+/- heterozygote mice. **c.** Head-twitch response for all three Cntnap2 genotype groups. Mean ± SEM.

**Extended Data Figure 2.**
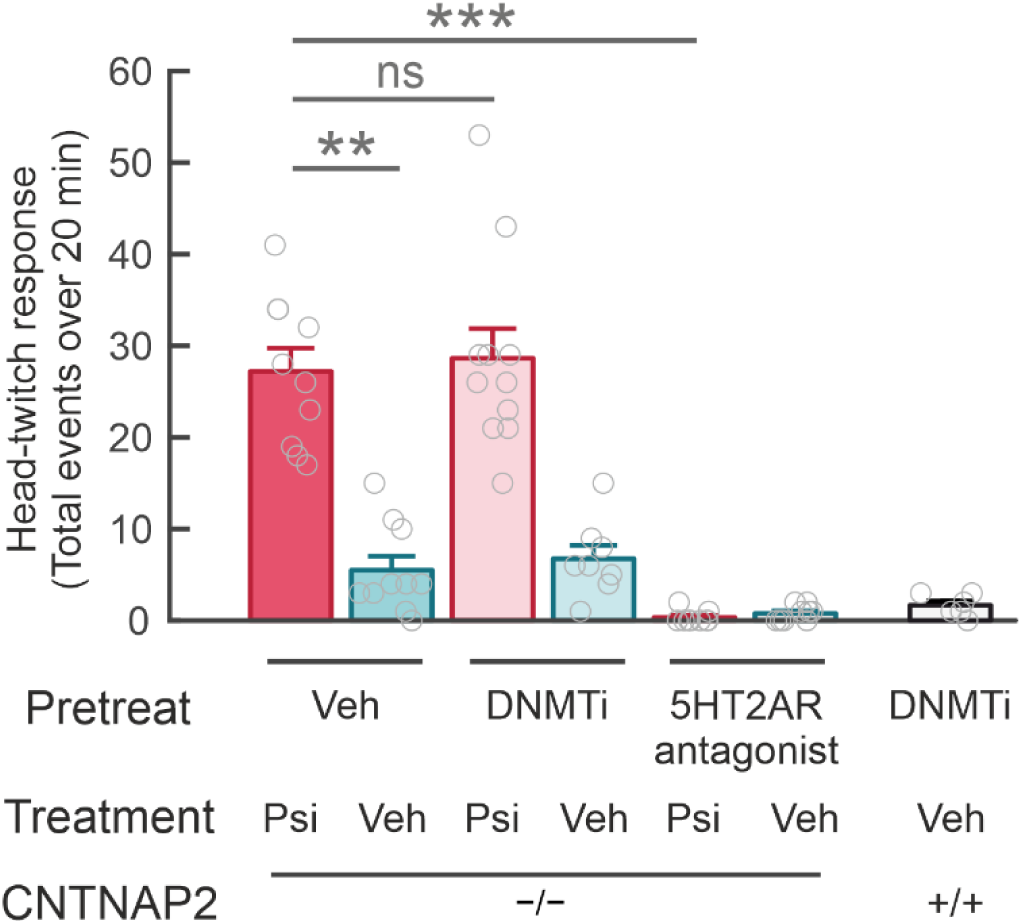
Head-twitch response for pretreatment experiments. Mean ± SEM.

**Extended Data Table 1.** Statistics for durability of psilocybin-induced effects on sociability, relates to Fig 1.

|  |  | Comparison group | Test used | Main effect | P value | Degrees of freedom |
| --- | --- | --- | --- | --- | --- | --- |
| **Extended Data Table 1a. Effect of psilocybin treatment on sociability in Cntnap2(-/-) compared to Cntnap2(+/+) mice** |  |  |  |  |  |  |
| Fig. 1a | Sociability index |  | Repeated measure ANOVA | Group | <0.001 | 3, 44 |
|  |  |  |  | Time | <0.001 | 2, 88 |
|  |  | Cntnap2(+/+) & Vehicle vs Cntnap2(-/-) & Vehicle | Bonferroni's post-hoc |  | 0.042 |  |
|  |  | Cntnap2(-/-) & Vehicle vs Cntnap2(-/-) & Psilocybin | Bonferroni's post-hoc |  | <0.001 |  |
|  |  | Cntnap2(+/+) & Vehicle vs Cntnap2(-/-) & Psilocybin | Bonferroni's post-hoc |  | 0.796 |  |
|  |  | Cntnap2(+/+) & Vehicle vs Cntnap2(+/+) & Psilocybin | Bonferroni's post-hoc |  | 0.106 |  |
| **Extended Data Table 1b. Effect of psilocybin treatment on sociability in Cntnap2(+/-) compared to Cntnap2(+/+) mice** |  |  |  |  |  |  |
| Extended Data Fig. 1c | Sociability index |  | Repeated measure ANOVA | Group | 0.049 | 3, 30 |
|  |  |  |  | Time | 0.023 | 2, 60 |
|  |  | Cntnap2(+/+) & Vehicle vs Cntnap2(+/-) & Vehicle | Bonferroni's post-hoc |  | 1.000 |  |
|  |  | Cntnap2(+/-) & Vehicle vs Cntnap2(+/-) & Psilocybin | Bonferroni's post-hoc |  | 0.368 |  |
|  |  | Cntnap2(+/+) & Psilocybin vs Cntnap2(+/-) & Psilocybin | Bonferroni's post-hoc |  | 1.000 |  |

**Extended Data Table 2.** Statistics for durability of psilocybin-induced effects on sociability by test session, relates to Fig 1.

|  | Comparison group | N | Descriptive stats |  |  |  | Test used | P value |
| --- | --- | --- | --- | --- | --- | --- | --- | --- |
|  |  |  | Group | Mean | Median | SEM |  |  |
| Extended Data Table 2a. Sociability first tested at 2 weeks after treatment |  |  |  |  |  |  |  |  |
| Sociability index | Vehicle vs Psilocybin | 15 | Cntnap2(-/-) & Vehicle | 0.129 | 0.101 | 0.050 | Independent T test | 0.025 |
|  |  | 15 | Cntnap2(-/-) & Psilocybin | 0.328 | 0.260 | 0.079 |  |  |
| **Extended Data Table 2b. Sociability by test session** |  |  |  |  |  |  |  |  |
| Sociability index | **1 day:**<br>Cntnap2(-/-) & Vehicle vs<br>Cntnap2(-/-) & Psilocybin | 19 | Cntnap2(-/-) & Vehicle | 0.374 | 0.380 | 0.058 | Independent T test | 0.001 |
|  |  | 15 | Cntnap2(-/-) & Psilocybin | 0.598 | 0.650 | 0.033 |  |  |
|  | **7 days:**<br>Cntnap2(-/-) & Vehicle vs<br>Cntnap2(-/-) & Psilocybin | 19 | Cntnap2(-/-) & Vehicle | 0.105 | 0.200 | 0.062 | Independent T test | <0.001 |
|  |  | 15 | Cntnap2(-/-) & Psilocybin | 0.498 | 0.500 | 0.053 |  |  |
|  | **14 day:**<br>Cntnap2(-/-) & Vehicle vs<br>Cntnap2(-/-) & Psilocybin | 15 | Cntnap2(-/-) & Vehicle | 0.028 | 0.100 | 0.104 | Independent T test | 0.028 |
|  |  | 15 | Cntnap2(-/-) & Psilocybin | 0.291 | 0.380 | 0.080 |  |  |
|  | **1 day:**<br>Cntnap2(+/+) & Vehicle vs<br>Cntnap2(+/+) & Psilocybin | 15 | Cntnap2(+/+) & Vehicle | 0.497 | 0.510 | 0.041 | Independent T test | 0.001 |
|  |  | 7 | Cntnap2(+/+) & Psilocybin | 0.151 | 0.130 | 0.076 |  |  |
|  | **7 days:**<br>Cntnap2(+/+) & Vehicle vs<br>Cntnap2(+/+) & Psilocybin | 15 | Cntnap2(+/+) & Vehicle | 0.331 | 0.260 | 0.080 | Independent T test | 0.107 |
|  |  | 7 | Cntnap2(+/+) & Psilocybin | 0.129 | 0.020 | 0.131 |  |  |
|  | **14 days:**<br>Cntnap2(+/+) & Vehicle vs<br>Cntnap2(+/+) & Psilocybin | 10 | Cntnap2(+/+) & Vehicle | 0.256 | 0.340 | 0.093 | Independent T test | 0.207 |
|  |  | 7 | Cntnap2(+/+) & Psilocybin | 0.149 | 0.140 | 0.088 |  |  |

**Extended Data Table 3.** Statistics for repetitive behaviour and locomotion, relates to Fig 1.

|  |  | Test session | Comparison group |  | Test used | N | Descriptive stats |  |  | P value |
| --- | --- | --- | --- | --- | --- | --- | --- | --- | --- | --- |
|  |  |  | Genotype | Treatment |  |  | Mean | Median | SEM |  |
| Extended Data Table 3a. Repetitive behaviour |  |  |  |  |  |  |  |  |  |  |
| Fig. 1c | Self-grooming episodes | Before treatment | Cntnap2(+/+) | - | Mann Whitney | 14 | 11.214 | 10.500 | 1.668 | 0.002 |
|  |  |  | Cntnap2(-/-) |  |  | 11 | 25.909 | 28.000 | 3.843 |  |
| Fig. 1d | Self-grooming episodes | 1 day after treatment | Cntnap2(-/-) | Vehicle | Mann Whitney | 5 | 30.200 | 26.000 | 2.577 | 0.329 |
|  |  |  | Cntnap2(-/-) | Psilocybin |  | 7 | 33.857 | 31.000 | 3.284 |  |
|  |  | 7 days after treatment | Cntnap2(-/-) | Vehicle | Mann Whitney | 6 | 30.833 | 26.000 | 4.246 | 0.461 |
|  |  |  | Cntnap2(-/-) | Psilocybin |  | 7 | 29.714 | 27.000 | 5.272 |  |
| Extended Data Table 3b. Locomotion |  |  |  |  |  |  |  |  |  |  |
| Fig. 1d | Distance travelled | Before treatment | Cntnap2(+/+) | - | Kruskal-Wallis with Dunn's post-hoc | 15 | 29200 | 28853 | 1737 | 0.001 |
|  |  |  | Cntnap2(-/-) |  |  | 22 | 38099 | 38158 | 1294 |  |
|  |  | Before treatment | Cntnap2(+/+) |  |  | 15 | 29200 | 28853 | 1737 | >0.999 |
|  |  |  | Cntnap2(+/-) |  |  | 10 | 32375 | 33378 | 2265 |  |
| Fig. 1e | Distance travelled | 1 day after treatment | Cntnap2(-/-) | Vehicle | Mann Whitney | 11 | 38583 | 39653 | 1095 | 0.518 |
|  |  |  | Cntnap2(-/-) | Psilocybin |  | 12 | 36990 | 37014 | 1620 |  |
|  |  | 7 days after treatment | Cntnap2(-/-) | Vehicle | Mann Whitney | 11 | 39930 | 41439 | 1526 | 0.976 |
|  |  |  | Cntnap2(-/-) | Psilocybin |  | 12 | 40714 | 38568 | 2069 |  |

**Extended Data Table 4.** Statistics for head-twitch response, and HTR-SI correlations.

Statistics for head-twitch response, and HTR-SI correlations.
|  |  | Comparison group |  | Test used | N | Descriptive stats |  |  | P value /<br>Adjusted P value |
| --- | --- | --- | --- | --- | --- | --- | --- | --- | --- |
|  |  | Genotype | Treatment |  |  | Mean | Median | SEM |  |
| Extended Data Table 4a. Head twitch response |  |  |  |  |  |  |  |  |  |
| Extended Data<br>Fig. 1d | HTR |  |  | Kruskal-Wallis |  |  |  |  | <0.001 |
|  |  | Cntnap2(+/+) | Vehicle | Dunn's multiple<br>comparison | 9 | 2.444 | 2.000 | 0.341 | 0.825 |
|  |  | Cntnap2(-/-) | Vehicle |  | 15 | 6.067 | 6.000 | 0.740 |  |
|  |  | Cntnap2(+/+) | Vehicle | Dunn's multiple<br>comparison | 9 | 2.444 | 2.000 | 0.341 | >0.999 |
|  |  | Cntnap2(+/-) | Vehicle |  | 6 | 3.000 | 2.000 | 1.072 |  |
|  |  | Cntnap2(+/+) | Vehicle | Dunn's multiple<br>comparison | 9 | 2.444 | 2.000 | 0.341 | <0.001 |
|  |  | Cntnap2(+/+) | Psilocybin |  | 7 | 22.429 | 23.000 | 1.462 |  |
|  |  | Cntnap2(-/-) | Vehicle | Dunn's multiple<br>comparison | 15 | 6.067 | 6.000 | 0.740 | 0.001 |
|  |  | Cntnap2(-/-) | Psilocybin |  | 15 | 21.267 | 22.000 | 1.300 |  |
|  |  | Cntnap2(+/-) | Vehicle | Dunn's multiple<br>comparison | 6 | 3.000 | 2.000 | 1.072 | 0.006 |
|  |  | Cntnap2(+/-) | Psilocybin |  | 6 | 19.500 | 18.500 | 2.332 |  |
|  |  | Cntnap2(+/+) | Psilocybin | Dunn's multiple<br>comparison | 7 | 22.429 | 23.000 | 1.462 | >0.999 |
|  |  | Cntnap2(-/-) | Psilocybin |  | 15 | 21.267 | 22.000 | 1.300 |  |
|  |  | Cntnap2(+/+) | Psilocybin | Dunn's multiple<br>comparison | 7 | 22.429 | 23.000 | 1.462 | >0.999 |
|  |  | Cntnap2(+/-) | Psilocybin |  | 6 | 19.500 | 18.500 | 2.332 |  |
| Extended Data Table 4b. HTR by sex |  |  |  |  |  |  |  |  |  |
|  | Male | Cntnap2(-/-) | Psilocybin | Mann Whitney | 6 | 19.167 | 21.500 | 2.151 | 0.452 |
|  | Female | Cntnap2(-/-) | Psilocybin |  | 9 | 22.667 | 22.000 | 1.546 |  |

| Extended Data Table 4c. HTR-SI correlations |  |  |  |  |  |  |  |
| --- | --- | --- | --- | --- | --- | --- | --- |
|  | 1 day after treatment | Cntnap2(+/+) | Psilocybin | Spearman's rho |  | rho = -0.655 | 0.110 |
|  |  | Cntnap2(-/-) | Psilocybin |  |  | rho = -0.397 | 0.143 |
|  |  | Cntnap2(+/-) | Psilocybin |  |  | rho = 0.714 | 0.111 |
|  | 1 week after treatment | Cntnap2(+/+) | Psilocybin | Spearman's rho |  | rho = -0.699 | 0.081 |
|  |  | Cntnap2(-/-) | Psilocybin |  |  | rho = 0.085 | 0.763 |
|  |  | Cntnap2(+/-) | Psilocybin |  |  | rho = 0.083 | 0.880 |
|  | 2 weeks after treatment | Cntnap2(+/+) | Psilocybin | Spearman's rho |  | rho = -0.538 | 0.213 |
|  |  | Cntnap2(-/-) | Psilocybin |  |  | rho = 0.084 | 0.766 |
|  |  | Cntnap2(+/-) | Psilocybin |  |  | rho = 0.421 | 0.406 |

**Extended Data Table 5.** Statistics for pretreatment on psilocybin’s effect on sociability, relates to Fig 2.

|  |  | Comparison group |  | Comparison group | Test used | Main effect | P value | Degrees of freedom |
| --- | --- | --- | --- | --- | --- | --- | --- | --- |
|  |  | Genotype | Pretreatment |  |  |  |  |  |
| Extended Data Table 5. Effect of pretreatment on psilocybin's effect on sociability in Cntnap2(-/-) mice |  |  |  |  |  |  |  |  |
| Fig. 2 | Sociability index |  |  |  | Repeated measure ANOVA | Group | <0.001 | 6, 57 |
|  |  |  |  |  |  | Time | 0.001 | 1, 57 |
|  |  | Cntnap2(-/-) | Vehicle | Vehicle | Bonferroni's post-hoc |  | <0.001 |  |
|  |  | Cntnap2(-/-) | Vehicle | Psilocybin |  |  |  |  |
|  |  | Cntnap2(-/-) | Vehicle | Psilocybin | Bonferroni's post-hoc |  | 0.003 |  |
|  |  | Cntnap2(-/-) | DNMTi | Psilocybin |  |  |  |  |
|  |  | Cntnap2(-/-) | Vehicle | Psilocybin | Bonferroni's post-hoc |  | 0.001 |  |
|  |  | Cntnap2(-/-) | 5HT2AR-ant | Psilocybin |  |  |  |  |
|  |  | Cntnap2(-/-) | DNMTi | Psilocybin | Bonferroni's post-hoc |  | 1.000 |  |
|  |  | Cntnap2(-/-) | DNMTi | Vehicle |  |  |  |  |
|  |  | Cntnap2(-/-) | Vehicle | Vehicle | Bonferroni's post-hoc |  | 1.000 |  |
|  |  | Cntnap2(-/-) | DNMTi | Vehicle |  |  |  |  |
|  |  | Cntnap2(-/-) | Vehicle | Vehicle | Bonferroni's post-hoc |  | 1.000 |  |
|  |  | Cntnap2(-/-) | 5HT2AR-ant | Vehicle |  |  |  |  |
|  |  | Cntnap2(-/-) | DNMTi | Psilocybin | Bonferroni's post-hoc |  | 1.000 |  |
|  |  | Cntnap2(-/-) | 5HT2AR-ant | Psilocybin |  |  |  |  |

**Extended Data Table 6.** Statistics for pretreatment on psilocybin’s sociability effect by test session, relates to Fig 2.

|  |  |  | Comparison group |  |  | N | Descriptive stats |  |  | Test used | P value |
| --- | --- | --- | --- | --- | --- | --- | --- | --- | --- | --- | --- |
|  |  |  | Genotype | Pretreatment | Treatment |  | Mean | Median | SEM |  |  |
| Extended Data Table 6. Sociability by test session |  |  |  |  |  |  |  |  |  |  |  |
| Fig. 2 | Sociability index | 1 day post-treatment | Cntnap2(-/-) | Vehicle | Vehicle | 10 | 0.207 | 0.220 | 0.038 | Independent T test | <0.001 |
|  |  |  |  | Vehicle | Psilocybin | 10 | 0.514 | 0.530 | 0.060 |  |  |
|  |  |  |  | Vehicle | Psilocybin | 10 | 0.514 | 0.530 | 0.060 | Independent T test | 0.005 |
|  |  |  |  | DNMTi | Psilocybin | 11 | 0.258 | 0.290 | 0.065 |  |  |
|  |  |  |  | Vehicle | Psilocybin | 10 | 0.514 | 0.530 | 0.060 | Independent T test | <0.001 |
|  |  |  |  | 5HT2AR-ant | Psilocybin | 9 | 0.177 | 0.210 | 0.059 |  |  |
|  |  |  |  | DNMTi | Psilocybin | 11 | 0.258 | 0.290 | 0.065 | Independent T test | 0.450 |
|  |  |  |  | DNMTi | Vehicle | 8 | 0.318 | 0.330 | 0.041 |  |  |
|  |  |  |  | Vehicle | Vehicle | 10 | 0.207 | 0.220 | 0.038 | Independent T test | 0.067 |
|  |  |  |  | DNMTi | Vehicle | 8 | 0.318 | 0.330 | 0.041 |  |  |
|  |  |  |  | Vehicle | Vehicle | 10 | 0.207 | 0.220 | 0.038 | Independent T test | 0.432 |
|  |  |  |  | 5HT2AR-ant | Vehicle | 9 | 0.251 | 0.230 | 0.039 |  |  |
|  |  |  |  | DNMTi | Psilocybin | 11 | 0.258 | 0.290 | 0.065 | Independent T test | 0.362 |
|  |  |  |  | 5HT2AR-ant | Psilocybin | 9 | 0.177 | 0.210 | 0.059 |  |  |
|  |  | 7 days post-treatment | Cntnap2(-/-) | Vehicle | Vehicle | 10 | 0.077 | 0.110 | 0.066 | Independent T test | 0.001 |
|  |  |  |  | Vehicle | Psilocybin | 10 | 0.475 | 0.525 | 0.090 |  |  |
|  |  |  |  | Vehicle | Psilocybin | 10 | 0.475 | 0.525 | 0.090 | Independent T test | 0.008 |
|  |  |  |  | DNMTi | Psilocybin | 11 | 0.155 | 0.140 | 0.080 |  |  |
|  |  |  |  | Vehicle | Psilocybin | 10 | 0.475 | 0.525 | 0.090 | Independent T test | 0.007 |
|  |  |  |  | 5HT2AR-ant | Psilocybin | 9 | 0.166 | 0.190 | 0.069 |  |  |
|  |  |  |  | DNMTi | Psilocybin | 11 | 0.155 | 0.140 | 0.080 | Independent T test | 0.926 |
|  |  |  |  | DNMTi | Vehicle | 8 | 0.144 | 0.155 | 0.082 |  |  |
|  |  |  |  | Vehicle | Vehicle | 10 | 0.077 | 0.110 | 0.066 | Independent T test | 0.537 |
|  |  |  |  | DNMTi | Vehicle | 8 | 0.144 | 0.155 | 0.082 |  |  |
|  |  |  |  | Vehicle | Vehicle | 10 | 0.077 | 0.110 | 0.066 | Independent T test | 0.435 |
|  |  |  |  | 5HT2AR-ant | Vehicle | 9 | 0.143 | 0.100 | 0.050 |  |  |
|  |  |  |  | DNMTi | Psilocybin | 11 | 0.155 | 0.140 | 0.080 | Independent T test | 0.918 |
|  |  |  |  | 5HT2AR-ant | Psilocybin | 9 | 0.166 | 0.190 | 0.069 |  |  |

**Extended Data Table 7.**
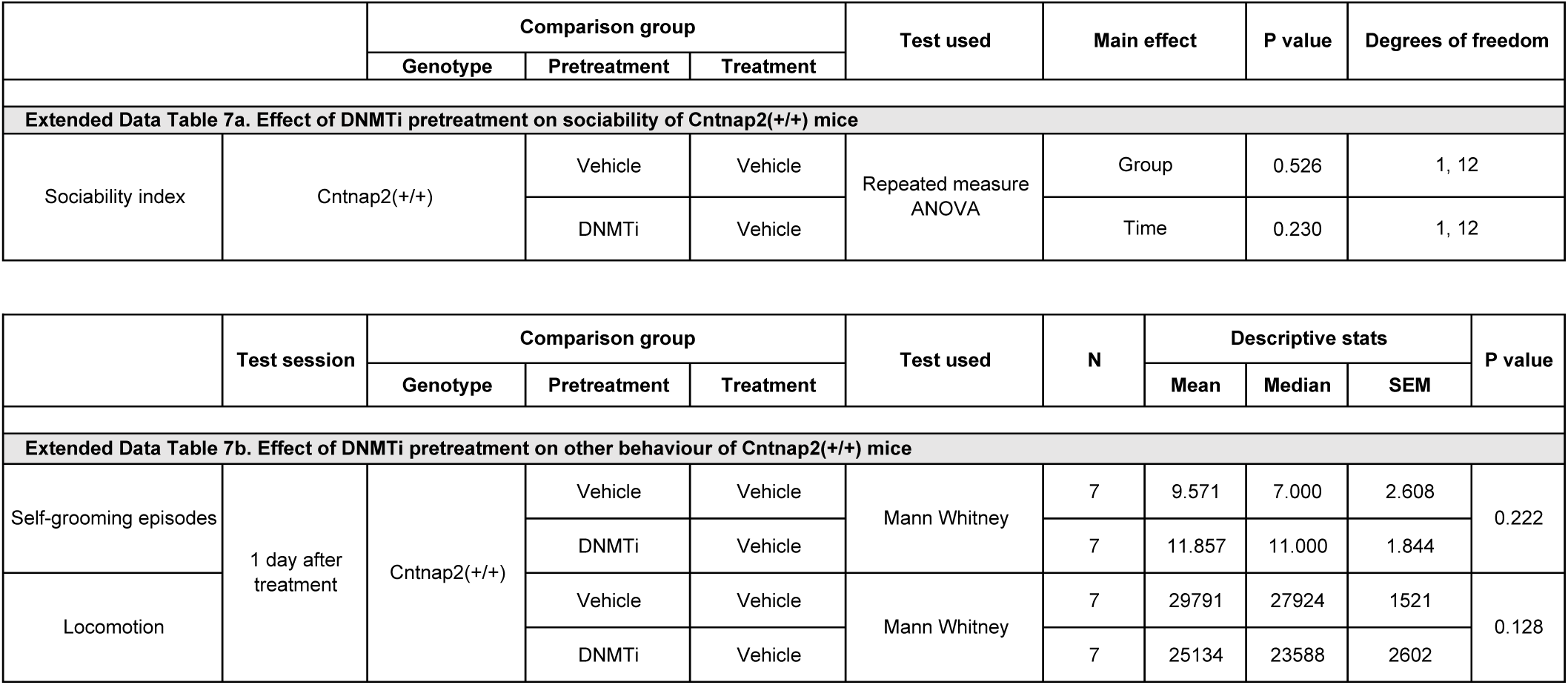
Statistics for the effects of DNMTi on behaviour of Cntnap2(+/+) mice.

**Extended Data Table 8.** Statistics for either DNMTi or 5HT2AR antagonist pretreatment on psilocybin’s effect on HTR.

|  |  | Comparison group |  |  | Test used | N | Descriptive stats |  |  | Adjusted P value |
| --- | --- | --- | --- | --- | --- | --- | --- | --- | --- | --- |
|  |  | Genotype | Pretreatment | Treatment |  |  | Mean | Median | SEM |  |
| Extended Data Table 8. Head twitch response |  |  |  |  |  |  |  |  |  |  |
| Extended Data Fig. 2 | HTR |  |  |  | Kruskal-Wallis |  |  |  |  | <0.001 |
|  |  | Cntnap2(-/-) | Vehicle | Vehicle | Dunn's multiple comparison | 10 | 5.500 | 4.000 | 1.529 | 0.0208 |
|  |  | Cntnap2(-/-) | Vehicle | Psilocybin |  | 10 | 27.200 | 27.000 | 2.542 |  |
|  |  | Cntnap2(-/-) | Vehicle | Psilocybin | Dunn's multiple comparison | 10 | 27.200 | 27.000 | 2.542 | <0.001 |
|  |  | Cntnap2(-/-) | 5HT2AR-ant | Psilocybin |  | 9 | 0.333 | 0.000 | 0.236 |  |
|  |  | Cntnap2(-/-) | Vehicle | Psilocybin | Dunn's multiple comparison | 10 | 27.200 | 27.000 | 2.542 | >0.999 |
|  |  | Cntnap2(-/-) | DNMTi | Psilocybin |  | 11 | 28.636 | 26.000 | 3.231 |  |
|  |  | Cntnap2(-/-) | Vehicle | Vehicle | Dunn's multiple comparison | 10 | 5.500 | 4.000 | 1.529 | 0.335 |
|  |  | Cntnap2(-/-) | 5HT2AR-ant | Vehicle |  | 9 | 0.778 | 1.000 | 0.278 |  |
|  |  | Cntnap2(-/-) | Vehicle | Vehicle | Dunn's multiple comparison | 10 | 5.500 | 4.000 | 1.529 | >0.999 |
|  |  | Cntnap2(-/-) | DNMTi | Vehicle |  | 8 | 6.750 | 6.000 | 1.461 |  |

## Notes

### Competing Interest Statement

The authors have declared no competing interest.

